# Conversation in fMRI: DMN Activates During Live Mentalizing

**DOI:** 10.64898/2026.09.07.749824

**Authors:** Ekaterina Torubarova, Caroline Arvidsson, André Pereira, Julia Uddén

**Affiliations:** KTH Royal Institute of Technology, Stockholm, Sweden; Stockholm University, Stockholm, Sweden

**Keywords:** Interaction, fMRI, DMN, FPN, Conversation, Canonical Networks

## Abstract

Although language might have evolved in a conversational setting, neural language processing is rarely studied in this setting. It is also important to delineate the roles of networks interacting with the language network, such as the default mode network and networks subserving social cognition or cognitive control. We studied the fundamental question of how conversational production and comprehension processes may differ. By acquiring fMRI data of participants engaging in conversations on ethical dilemmas with a confederate, we aimed to study extra-linguistic areas during live mentalizing, crucially with long (10 min) conversations. A subject-specific network analysis was used to complement group-averaged whole-brain analysis. The results replicate asymmetries between production and comprehension already observed using short conversations. This included left inferior frontal gyrus preferences for production and superior and middle temporal gyri preferences for comprehension. Activation in the posterior dorsal medial prefrontal cortex was specific to production, a result that is stable across conversational durations. Notably, we found preferential activity of the default mode network (DMN) during comprehension, specifically during longer conversations. The frontoparietal network for control (FPN) showed the same pattern, although it was less robust. This might indicate simultaneous self-reflective, Theory of Mind, and/or situational processing during comprehension, potentially due to lower conversational pressures, compared to production. The subnetwork DN-B was more strongly activated than DNA, across both comprehension and production, suggesting that Theory of Mind remains continuously active throughout conversation. Our findings thus advance the understanding of how several networks contribute to production vs comprehension during conversation.

## Introduction

Let us suppose that it is true that language (here meaning the externalization of cognition via speech, sign, or writing (Fitch, 2017)) evolved for communication (Fedorenko et al., 2024), or interaction (Levinson, 2019), rather than thought itself (for a review, see Fitch, 2017). Then, it follows that conversation is the benchmark niche when studying language processing experimentally (Levinson, 2024), as key processes crucial for context-relevant production and comprehension will be largely absent in non-conversational settings. In conversational production, for instance, we shape our view on a topic and express it to engage the listener. In conversational comprehension, we interpret words and gestures in context and anticipate where the interlocutor aims to steer the conversation. Pressure for timing in conversation further obliges both participants to “analyze each utterance across its delivery” (Sacks et al., 1974; Bögels and Levinson, 2017). Indeed, there is a movement in the neurobiology of language to collect data in naturalistic conditions (Matusz et al., 2019; Hamilton and Huth, 2020), as opposed to using common isolation paradigms (called “language in vacuum” paradigms by their critics, e.g. Clark, 1997). Simultaneously with this movement, another trend is getting prominent regarding the functional anatomical focus in the field, with additional regions and networks beyond the core language network being considered when studying language (Hagoort, 2014, 2019; Hickok, 2022; Fedorenko et al., 2024). The current paper aligns with both of these movements, using the novel approach of taking the full range of processing in the linguistic and extra-linguistic networks during conversation as its explanandum. In the following sections, we will discuss the extralinguistic networks likely to be involved during conversation, and whether they are engaged differently in production and comprehension.

### Possible Extralinguistic Networks Involved During Conversation

In contemporary cognitive neuroscience, canonical brain networks that show functional connectivity, even at rest, are pivotal building blocks for uncovering the function of the mind (see, for instance, Du et al., 2024). While the language network (see LANG in Fig. 1) activates for language-in-isolation, at least two additional such networks are expected to contribute during conversation: the default mode network (DMN; Du et al., 2024) and the fronto-parietal network (FPN; Du et al., 2024; in other works also called multiple demand network: MD; Duncan, 2010).

**Figure 1.**
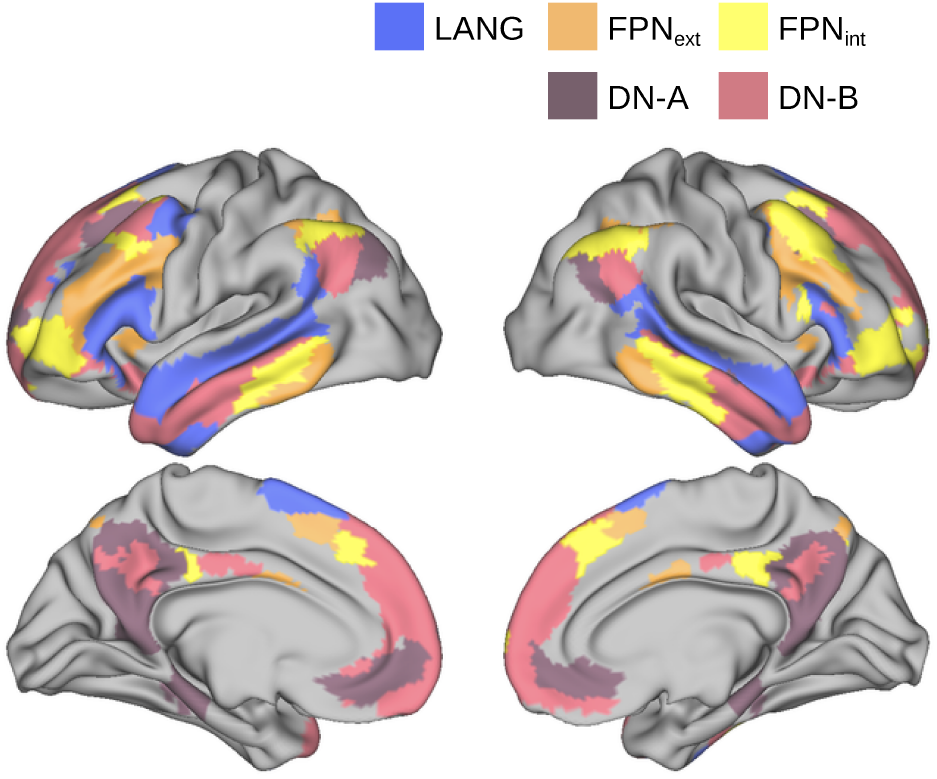
Canonical networks from Du et al. (2024) Consensus atlas, projected onto fs_LR 32k mid-thickness surface. For associating network labels with their established functions, we use the following references for the Du et al. (2024) atlas: FPN-A - FPN_ext_; FPN-B - FPN_int_.

The DMN comprises a set of regions in the medial pre-frontal cortex, the posterior cingulate cortex, and lateral parietal and temporal cortices (Fig. 1). While initially suggested to represent brain at rest (Raichle et al., 2001), later these regions were found to show high activity during internally focused processing (e.g., self-generated thought), and reduced activity during externally oriented, goal-directed tasks (Andrews-Hanna et al., 2014). An early group-average clustering approach suggested a division of DMN into two subnetworks: a dorsomedial network for mentalizing and Theory-of-Mind (involving the dorsomedial prefrontal cortex (dmPFC), temporo-parietal junction (TPJ), lateral temporal cortex, temporal pole, and inferior frontal gyrus (Andrews-Hanna et al., 2010)), and a medial temporal network for personal situational modeling in time and space (featuring the hippocampus, parahippocampal cortex, retrosplenial cortex, and posterior inferior parietal lobe (Andrews-Hanna et al., 2010)); each interacting with a common core (involving the anterior medial prefrontal cortex (amPFC), posterior cingulate cortex (PCC), bilateral angular gyrus, lateral temporal lobes, and superior frontal gyrus (Andrews-Hanna et al., 2010)), serving as an integration hub (Andrews-Hanna et al., 2010; Spreng and Andrews-Hanna, 2015; Yeo et al., 2011). More recent within-individual precision mapping instead identifies two parallel subnetworks: DN-A and DN-B, with the closely juxtaposed “core” regions belonging to one or the other of these networks (Braga and Buckner, 2017; Du et al., 2024). The function of DN-A has been linked to internal situational/episodic remembering and spatiotemporal projection; while DN-B has been linked to mentalizing and Theory of Mind (DiNicola and Buckner, 2021), thus roughly corresponding to the previous medial-temporal and dorsomedial default network subsystems, respectively. Related suggestions are that the two DMN subnetworks collectively encode *situation models* (Fernandino and Binder, 2024; Thye et al., 2024b) and/or mental state and intention inference, for self and others (Andrews-Hanna et al., 2010; for a precise representational suggestion, see Thornton and Tamir, 2024).

Situation models are experience-based mental representations of events in time and space, in other words “the state of affairs”. They encode the entities involved, as well as their properties and relationships, capturing the situation as *implied* by the input, rather than its immediate perceptual or semantic features (Zwaan and Radvansky, 1998; Barnett and Bellana, 2025). Importantly, considering that the participants of a conversation are required to construct shared situation models and infer mental states, these suggestions predict that the DMN is involved during conversation (as supported by initial recent evidence in Spatola and Chaminade, 2022 and Yamashita et al., 2025).

The function of the FPN is, in short, cognitive control (e.g., enabling flexible planning and regulation of complex tasks, including working memory tasks, as well as novel higher-order tasks). Spatially, FPN is interdigitated with DMN (Fig. 1) and follows a similar organizational motif, with regions distributed in frontal, parietal, and temporal cortices (Braga et al., 2020). Some non-conversational comprehension studies suggest a dissociation between the language network and FPN (e.g., Paunov et al., 2019; Blank et al., 2014; Schieche et al., 2025). Fedorenko et al. (2024), however, suggest involvement of FPN in language processing during, e.g., effortful comprehension or processing of logical statements. One can thus anticipate that involvement of FPN might depend on the processing costs that context incurs on the listener. Developmental studies also highlight the importance of executive functions for interactive language use (Matthews et al., 2018).

Like DMN, the FPN has been delineated into two parallel networks: FPN-A and FPN-B (Braga et al., 2020; Du et al., 2024). Anatomically, FPN-A and FPN-B both involve a heterogenous set of regions across the lateral inferior frontal cortex, lateral temporal and parietal regions around the intraparietal sulcus. Importantly however, which of the two networks that is named FPN-A and FPN-B has differed across studies, as we will see. To distinguish these subnetworks, one can rely on their spatial topography and connectivity profiles (Dixon et al., 2018; Braga et al., 2020): one FPN network (linked to DMN) typically lies more ventral in inferior parietal cortex (neighboring DN-B) and more anterior in lateral temporal cortex, and is functionally associated with internally-oriented stimulus-independent executive control; while the other (linked to the dorsal attention network, DAN) is typically more posterior and dorsal, respectively, and is associated with attentional control to task-relevant stimuli and working memory demands. Both networks however support domain-general control (Dixon et al., 2018; Du et al., 2024). In Braga et al. (2020), the DMN-linked ventral/anterior internal control network is named FPN-A, and the DAN-linked dorsal/posterior external control network is called FPN-B, while in the frequently used Du et al. (2024) and Yeo et al. (2011) parcellation atlases, the labels are the opposite (FPN-B/FPN-A and ContB/ContA, respectively). To resolve the ambiguity, throughout the paper we will refer to these networks as FPN_int_ and FPN_ext_, to reflect their functional asymmetry in terms of internal/external control orientation.

### Possible Differential Involvement Of Extralinguistic Networks In Production Vs Comprehension

Fedorenko et al. (2024) sketch the information flow between the DMN, FPN, and language networks in comprehension and production. Comprehension starts in auditory areas and flows via the language network (which is crucially a network for “meaning dressed in language”, see Fedorenko et al., 2024) to these extralinguistic networks. Production, on the other hand, starts in the extralinguistic networks (named “higher-level knowledge and reasoning areas” by the authors of Fedorenko et al., 2024), via the language network, down to more peripheral motor planning and motor areas. The DMN is at the top of the hierarchy of functional networks, with the language and FPN networks in the middle, and sensory-motor networks at the lower end (Margulies et al., 2016). Overlaying this hierarchy onto the processing flow proposed by Fedorenko et al. (2024), comprehension appears to be more “bottom-up”, moving from the language network to the midline areas, including DMN and FPN (although there is also predictive processing during comprehension, Kuperberg and Jaeger, 2016). Production is relatively more “top-down” with respect to this global hierarchy between networks, as it moves in the opposite direction.

Our previous study (Arvidsson et al., 2024b) is one of just a few studies that test such a model within one study, in a conversational setting. In that study, we used the conversational situation to get within-participant comparisons of production and comprehension in fMRI. We found partial overlap of these two systems in the language network via a conjunction analysis of production/comprehension vs fixation. However, activation of some regions was specific to either production or comprehension, as revealed by directly contrasting production and comprehension. For instance, the left inferior frontal gyrus (LIFG) was preferentially active for production, as also observed in a naturalistic monologue vs listening paradigm (Giglio et al., 2024). We also identified stronger activity for production than comprehension in the midline part of the superior frontal gyrus, including the supplementary motor area (SMA). While this area has traditionally been understood as subserving motor processes (e.g., Tanji, 1994; Boy et al., 2010) and is typically overlooked in large-scale models of the neurobiology of language, there is accumulating evidence suggesting that the anterior aspect of this region, i.e. the preSMA, subserves higher-order language processes (Alario et al., 2006; Wolna et al., 2026; DiNicola and Buckner, 2021; for a review, see Hertrich et al., 2016). Furthermore, activity in the precentral gyrus was specific to production in Arvidsson et al. (2024b), a finding which replicated and was emphasized as higher-order and more linguistic by Goldstein et al. (2025), who used conversational data recorded with electrocorticography (ECoG). Together, these results serve as empirical evidence supporting the idea of frontal higher-order areas showing early preferential activity for conversational production.

In Arvidsson et al. (2024b), we did not observe any significant extralinguistic network activity when contrasting comprehension to production (see possible reasons in the next paragraph). However, studies using naturalistic paradigms, albeit not conversational, have found extralinguistic processing in both production and comprehension. The ventromedial prefrontal cortex supported speech preparation in interactive versus non-interactive contexts (Kuhlen et al., 2017) and subserved producing and interpreting communicative signals as measured with MEG (Stolk et al., 2016). DMN regions (e.g., the temporal cortices, the posterior cingulate cortex, and the anterior medial prefrontal cortex) were also activated during narrative production and comprehension (Silbert et al., 2014; Stephens et al., 2010). While large-scale models of network interactions have been posed at the whole network level (as in Fedorenko et al., 2024), this empirical data altogether suggests the possibility of partial involvement of extralinguistic networks, both in comprehension and production.

Parameters such as the methods used to elicit the conversations, as well as the analytical approach, need to be considered. For instance, while Arvidsson et al. (2024b) used short 1-minute conversations and found no DMN-activity, a second study on the same data showed increased responses after several short conversations with the same agent in a posterior DMN-area, uniquely for human-human conversations (vs conversation with a robot, Spatola and Chaminade, 2022). A recent fMRI study by Yamashita et al. (2025) emphasizes the importance of pragmatic (compared to linguistic) signals during longer conversations and shows a role of the DMN network during conversation (see further specification in the discussion).

Moreover, most of the existing findings on DMN and/or FPN involvement in naturalistic language use network ROIs defined via group averaging. Such methods do not allow to differentiate between their relevant subnetworks, as a growing body of literature suggests that the boundaries between tightly interdigitated associative networks can be blurred and bleed into each other, when using a group average approach (Fedorenko et al., 2010; Braga and Buckner, 2017; Gordon et al., 2017; Kong et al., 2019; Shain and Fedorenko, 2026). We therefore use subnetworks that are defined considering individual subjects’ functional connectivity, to further specify the involvement of these networks during conversation.

### The Current Study

Given the evolutionary importance of conversation, together with the observation of radically different pressures on the speaker and listener (Levinson and Torreira, 2015; Levinson, 2024), we aimed to further study a potential dissociation between conversational production and comprehension, but now with longer conversations. In the current paper, we focus on whether and how the extra-linguistic DMN and FPN networks are involved in production and comprehension.

We have published a novel open participant-confederate conversational dataset (N = 30), to address previous data limitations (i.e. Arvidsson et al., 2024b), e.g. with regard to the short length of conversations (Torubarova et al., 2025). The paradigm builds on a previous dataset (Rauch-bauer et al., 2020), involving short 1-minute conversations with a confederate. We made the essential change to now elicit longer conversations by using more complex discussion topics. Participants partook in three 10-minute conversations, discussing their opinions on ethical dilemmas. This task crucially involves “live mentalizing” (Hamilton and Holler, 2023), as it involves managing both their own and their interlocutors’ beliefs and opinions and react to the changes in confederate’s behavior across conversations. This paradigm, while naturalistic, lends itself well to studying the role of extra-linguistic networks during language processing, since it requires planning and reflection on one’s and other’s viewpoint and relation to past personal experience. We use state-of-the-art network definitions based on the individual subject’s functional connectivity to answer our questions. Our main novel questions for the current paper are a) whether the DMN and/or FPN networks and their subnetworks DN-A, DN-B, FPN_ext_ and FPN_int_ are involved during live mentalizing in the form of conversation; b) are there discrepancies in the involvement of these networks across production vs comprehension.

## Materials And Methods

### Data

This study used an open-source dataset collected by the current authors (Torubarova et al., 2025). We only analyzed the human-human conversations, including 30 participants with normal or corrected vision and hearing, no reported neurological, psychiatric, or language-related disorders. All participants were native Swedish speakers (aged 21-39, mean = 26.87 y/o, 15 female). Participants engaged in three 10-minute-long unscripted conversations with the same 32 y.o. female confederate presented as a local student. Each conversation centered around a pre-defined ethical dilemma. These dilemmas were chosen to provide potentially engaging 10-minute conversations that could involve extralinguistic processes naturally. The task of the participants was to freely discuss the topics as if they were having a conversation with a friend. Across the three conversations, the level of engagement was manipulated via confederate’s verbal and nonverbal behavior, namely the number of backchannels (i. e. short responses serving to provide feedback or indicate attentive listening, such as ‘uhum’ or ‘a-ha’), reactivity in answering questions and asking for clarification, and proactivity in moving the conversation forward. This way, the confederate’s role varied from an engaged communicator to a passive listener, with an intermediary active listener role always fixed in the second conversation. This aspect of the design is however not the focus in the current paper. The participant and confederate were connected via a bidirectional audio and unidirectional video link. The participant could see the confederate on the screen in real time, but not vice versa. The opposite video link was ruled out due to the scanner coil blocking the participant’s face. Both audio and video streams were continuous throughout the whole conversation, allowing for spontaneous behavior from both the participant and the confederate. To mitigate the influence of the MRI scanner noise on the interaction, active noise cancellation was applied, and the participant’s voice was further denoised in real-time using external processing software.

### fMRI Data Acquisition and Preprocessing

The fMRI data consisted of one structural and three functional runs, consisting of 15 seconds of looking at a fixation cross, 10 minutes of conversation, and another 15 seconds of looking at the fixation cross. The images were acquired with a 3T 20-channel coil scanner with the following parameters for functional images: EPI sequence, TR 1.2 s, multiband factor 3.

The data were preprocessed in fMRIPrep v21.0.1 (Este-ban et al., 2018), including slice-timing correction, realignment, and co-registration with structural T1w images into MNI space. To account for head movement, SPM12 ArtRepair^1^ toolbox was used to repair volumes with >1mm framewise displacement (FD) by interpolation (see motion statistics in Table 1). While this is an unconventionally lenient motion threshold, we naturally assumed larger motion in our data compared to isolation studies; and we deliberately chose a higher interpolation threshold to preserve temporal continuity of the data. Full details on image acquisition and preprocessing steps are given in Torubarova et al. (2025).

**Table 1.** Descriptive statistics of the data per subject for each engagement condition: average number and duration of events, mean framewise displacement (FD) and percentage of scans with excessive motion (FD *>* 1mm). See statistics breakdown by engagement level in Supplementary Table 1.

| Event Type | N of Events | Event Duration (s) | Mean FD (mm) | FD > 1mm (scans) |
| --- | --- | --- | --- | --- |
| **Production** | 143.4 (SD = 27.2) | 2.02 (SD = 1.53) | 0.28 (SD = 0.26) | 2.35% (SD = 1.25) |
| **Comprehension** | 103.9 (SD = 23.0) | 2.12 (SD = 2.46) | 0.21 (SD = 0.21) | 1.28% (SD = 0.85) |
| **Silence** | 147.4 (SD = 28.1) | 0.90 (SD = 0.82) | 0.25 (SD = 0.22) | 1.50% (SD = 0.95) |

### Whole-Brain Analysis

To define the conversational events of interest, the conversations were automatically segmented into speech units surrounded by silence *≥*200 ms, and segmentations were manually checked to ensure their correctness. Based on this segmentation, for the current analysis, we defined: production (participant’s speech), comprehension (confederate’s speech), and silence (Table 1). In addition, we also defined turn initiation as a 600 ms time window before the onset of production. This event type was included in the first-level model, but its analysis was out of the scope of the current paper.

Events below 300 ms were removed from the model to ensure that the hemodynamic fMRI response was adequately captured. A general linear model (GLM) was used with five conversational events as regressors: production, comprehension, turn initiation, silence, and fixation cross. The GLM included six realignment parameters from motion correction (x, y, z translations and rotations), not convolved with the canonical HRF.

The following contrasts were created at the first level, following Arvidsson et al. (2024b): 1.a) production > fixation cross, 1.b) comprehension > fixation cross; 2.a) production > comprehension, 2.b) comprehension > production. The contrasts 1.a and 1.b provide a bird’ s-eye view of the regions involved in these processes, as studied individually, while contrasts 2.a and 2.b investigate the extent to which they share neural infrastructure or are different. While no smoothing was applied to BOLD time-series, individual first-level contrast maps were smoothed in SPM12 using a 6-mm Gaussian kernel.

The second-level group analysis was conducted with one-sample t-tests. To find overlap between contrasts 1.a and 1.b, we also performed a conjunction analysis as implemented in SPM12. For this analysis, we estimated another second-level model using a full factorial design with the system (production and comprehension) as a factor, using first-level contrast images. The model treated observations as dependent and assumed equal variance. Conjunctions were tested against the conjunction null (Nichols et al., 2005). To ensure that the group level overlap between contrasts 1.a and 1.b was not an artifact of spatial blurring, we also calculated the spatial overlap of individual subjects’ contrast maps for production > fixation and comprehension > fixation via Dice Similarity Coefficient (DSC), setting a threshold to the top 10% of voxels sorted by t-value for each contrast (Blank et al., 2016) to account for global signal variability.

Cluster-forming threshold was set to *P*_FWE_ < 0.05. Family-wise error correction, as implemented in SPM12, was used as the multiple comparison correction method (cluster and peak level). In the results, only clusters with *P*_FWE_ < 0.05 at the cluster level are reported. Anatomical labeling was done using the ROI MNI V7 anatomical parcellation database in the AAL3 toolbox in SPM12 (Rolls et al., 2020). Only regions occupying at least 10% of the cluster and clusters of at least 15 voxels were included in the report. All results are visualized on a fs_LR 32k mid-thickness surface using Connectome Workbench wb_-view platform, with only significant clusters shown.

### Subject-Specific Functional Network Analysis, Step 1: Estimation

We conducted a subject-specific network analysis, as a follow-up analysis suggested in the review processes, to: (1) explicitly test for network and subnetwork involvement in production and comprehension, and (2) to exclude the possibility that interdigitated networks bleed into each other as a group averaging artefact. This analysis complements the whole-brain results. To estimate subject-specific functional networks, we used the Multi-Session Hierarchical Bayesian Model (MSHBM, Kong et al., 2019) parcellation algorithm, which can reliably estimate 17 functional networks from a small amount of both resting-state and task-based data by using a population-level spatial prior. Alternative popular methods in dense precision mapping studies, e.g. seed-based or k-means parcellation, require a large amount of data and are susceptible to intra-subject noise at standard scan durations.

To fit into the MSHBM pipeline, we first projected the BOLD timeseries to fsaverage5 surface space using the Nilearn Python library. Then, to isolate background connectivity from task-evoked activity (Fair et al., 2007), we used a GLM to regress out (1) “task confounds”, which in this case are the potential confounds from the conversational events (production, comprehension, silence, fixation cross) convolved with the SPM canonical HRF, as well as (2) standard fMRIPrep confounds including 6 rigid body motion parameters, white matter (WM) and cerebrospinal fluid (CSF) signals. We then applied a high-pass filter at 0.008 Hz (default in SPM). In addition, we censored volumes with > 0.5 mm FD to account for excessive motion. Since MSHBM uses data from all runs of each subject, we used a stricter motion threshold than in the whole-brain analysis. To ensure a sufficient amount of data after preprocessing, we checked that across all three runs for each subject, the amount of data passing motion censoring summed to at least 5 minutes, which all subjects passed. We then ran MSHBM parcellation as implemented in the Yeo lab CBIG toolbox^2^.

To validate subject-specific network estimation, since the algorithm carries no information of the “ground truth” network label, we calculated a spatial overlap between the canonical atlas and subject-specific networks, taking the highest overlap as the closest network match. Since MSHBM uses the Yeo et al. (2011) atlas which does not feature a dedicated language network (this was one of our main interests), we additionally validated subject-specific networks via the Du et al. (2024) Consensus atlas. This atlas features a dedicated language network (labeled LANG), alongside two FPN and DMN subnetworks. The mapping between canonical and subject-specific networks was consistent for all subjects for both the Yeo et al. (2011) and Du et al. (2024) atlases. Importantly, the subject-specific language network candidates featured the core left-lateralized fronto-temporal regions, consistent with the well-established language network topography (Fedorenko and Thompson-Schill, 2014; Shain and Fedorenko, 2026). Moreover, for every subject-specific network, intra-subject stability across runs was greater than inter-subject overlap (*P* < 0.001), in accordance with Kong et al. (2019). In the Supplementary Material S-2 we provide: (1) an explicit test that validates the subject-specific language networks, (2) analysis showing greater intra-subject stability than inter-subject overlap, (3) a consensus atlas of all the subject-specific networks.

### Step 2: ROI Analysis

For the ROI analysis continuing from the creation of subject-specific networks in Step 1, we first projected every subject’s first-level contrast images to fsaverage5 surface space. Then, we defined seven ROIs derived from subject-specific network masks: DN-A, DN-B, FPN_ext_, FPN_int_, LANG, plus DMN (Overall) and FPN (Overall) defined as a spatial union of the respective subnetworks. We then extracted the mean contrast estimates across vertices within each of these ROIs for the contrasts production > fixation and comprehension > fixation. We used a series of one-sample t-tests to evaluate how strongly each network was involved in production/comprehension vs baseline. Paired t-tests were used for the comparison of each network between production and comprehension, and for the relative contribution of DN-A vs DN-B and FPN_ext_ vs FPN_int_ within production and comprehension, separately.

Moreover, to investigate previously observed dissociations within the frontal (left IFG) and temporal (left STS) nodes of the language network in conversational data (Arvidsson et al., 2024b), we identified these nodes within our subject-specific language network maps. To isolate these individualized functional nodes, we intersected each subject-specific language network with LIFG and LSTS ROIs from Fedorenko et al. (2010). To each ROI, we added 10 mm padding, to maximize the coverage of subject-specific maps variability. For each individually established ROI, we extracted mean contrast estimates for the production > fixation and comprehension > fixation. Paired t-tests were then conducted to evaluate the differential involvement of these nodes in production vs comprehension.

## Results

### Whole-Brain Analysis

Production and comprehension were contrasted against the fixation cross as an explicit baseline. The contrast **production > fixation** yielded 16 significant clusters (Fig. 2): (1) right precentral and postcentral gyri, superior and middle temporal gyri; (2) left precentral and postcentral gyri; (3, 11, 15) cerebellum; (4, 14) bilateral supplementary motor area; (5, 6) left middle and bilateral superior temporal gyrus; (7) left inferior frontal gyrus; (8-10) basal ganglia; (12) left superior temporal pole; (13, 16) bilateral calcarine sulcus, left lingual and middle occipital gyri.

**Figure 2.**
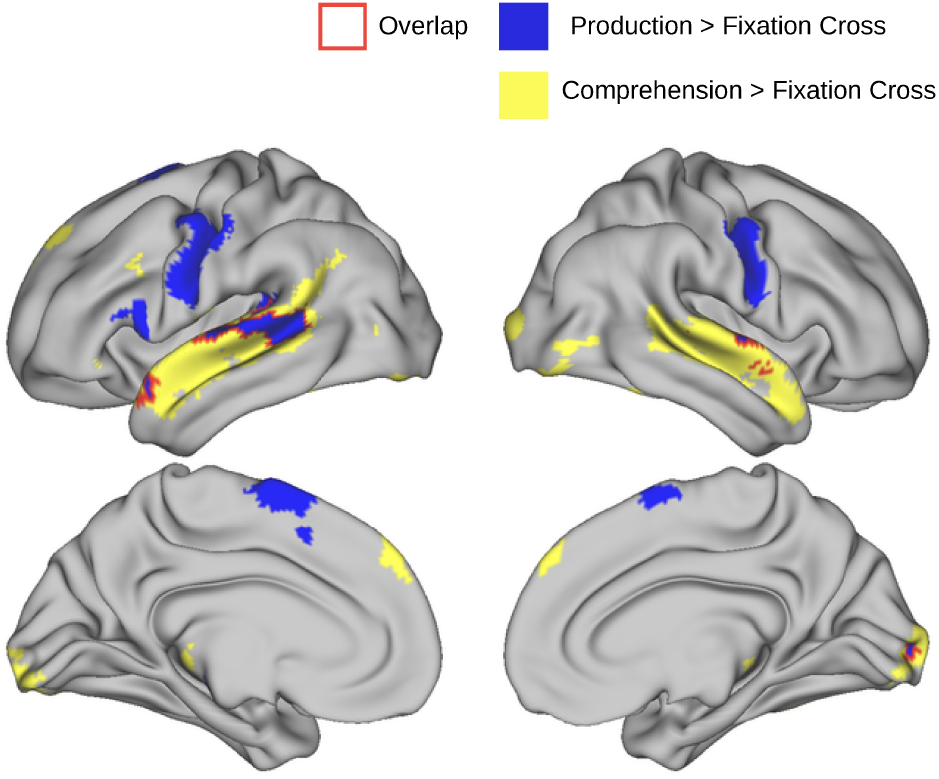
Whole-brain results for contrasts production vs fixation cross and comprehension vs fixation cross. The red outline shows the results of the conjunction analysis, limited to vertices overlapping between these contrasts. Note that a portion of overlap within the Sylvian fissure is not visible in this view. Cluster-forming threshold for these contrasts was set to *P* _FWE_ *<* 0.05 (no extend-level threshold, k = 0). In all figures, only significant clusters are shown (*P* _FWE_ *<* 0.05).

The contrast **comprehension > fixation** yielded 18 significant clusters (Fig. 2): (1-2) bilateral middle and superior temporal gyri; (3) bilateral calcarine sulcus; (4, 5, 9,

14, 17) cerebellum; (6, 12) bilateral fusiform and inferior temporal gyri; (7) right inferior occipital gyrus, right middle temporal gyrus; (8) bilateral superior frontal gyrus; (10, 11) left thalamus, left lingual gyrus; (13, 15) left inferior frontal gyrus; (16) left middle frontal and left precentral gyri; (18) left middle occipital gyrus.

A conjunction analysis showed overlap between these contrasts across 15 clusters (Fig. 2, note that only the voxels overlapping between production > fixation and comprehension > fixation are shown): (1) right superior temporal and right Heschl’s gyri; (2) left superior temporal pole; (3) left middle and superior temporal gyri; (4, 5, 14) cerebellum; (6) bilateral calcarine sulcus; (7-8) bilateral fusiform gyrus and cerebellum; (9, 13, 15) left inferior frontal gyrus; (10) right middle and superior temporal pole; (11-12) left superior frontal gyrus. See all significant clusters with their MNI coordinates in Supplementary Table 2. At the individual subject level, the average spatial overlap between production > fixation and comprehension > fixation contrasts had a Dice index of 0.55 (SD = 0.07).

The reverse contrast, **comprehension > production**, yielded increased activation in 20 clusters (Fig. 3): (1) bilateral precuneus and midcingulate cortex; (2, 6) left middle and superior temporal gyri; (3) right middle and superior temporal and angular gyri; (4) bilateral anterior cingulate cortex, left superior frontal and bilateral middle frontal gyri; (5, 16) bilateral hippocampus and parahippocampal gyri, left amygdala; (7) basal ganglia; (8-9) bilateral fusiform and inferior temporal gyri; (10) cerebellum; (11) left angular, middle temporal and middle occipital gyri; (12) left middle and inferior occipital gyri and left calcarine sulcus; (13) right lingual and inferior occipital gyri, right calcarine sulcus; (14-15, 18) bilateral orbitofrontal cortex and superior/ middle temporal pole, left insula; (17) right middle frontal gyrus; (19) bilateral superior frontal gyrus; (20) right inferior frontal gyrus.

### ROI Analysis

The results of ROI analysis are presented in Tables 2-4. DMN, FPN (when taken as a spatial union of the respective subnetworks) and the language network showed stronger activation in comprehension compared to production (DMN mean difference = 0.93, *T*_*29*_ = 7.29, *P* < 0.001; FPN mean diff. = 0.32, *T*_*29*_ = 2.95, *P* = 0.006; LANG mean diff. = 1.19, *T*_*29*_ = 7.29, *P* < 0.001). Additionally, the overall DMN and FPN showed a significantly stronger activation in comprehension compared to fixation cross baseline (*P* < 0.05), but null activation in production compared to the baseline (*P* > 0.05). When analyzing subnetworks individually, all four subnetworks showed a positive activation in comprehension compared to baseline (*P* < 0.05), with DN-B stronger than DN-A (mean diff. = 0.51, *T*_*29*_ = 5.6, *P* < 0.001) and FPN_ext_ not significantly different from FPN_int_ (mean diff. = -0.04, *T*_*29*_ = -0.75, *P* = 0.46). In production, DN-A showed a significantly lower activation than baseline (mean diff. = -0.86, *T*_*29*_ = -3.85, *P* < 0.001), and FPN_int_ showed slightly lower, but not significant activation (mean diff. = -0.22, *T*_*29*_ = -1.13, *P* = 0.26). On the contrary, DN-B showed a significantly higher activation compared to baseline (mean diff. = 0.71, *T*_*29*_ = 3.52, *P* < 0.01), and FPN_ext_ a slightly higher, but not significant activation (mean diff. = 0.2, *T*_*29*_ = 0.97, *P* = 0.33). For production, DN-B was relatively stronger than DN-A (mean diff. = 1.31, *T*_*29*_ = 12.4, *P* < 0.001), and FPN_ext_ stronger than FPN_int_ (mean diff. = 0.21, *T*_*29*_ = 2.63, *P* = 0.01).

**Table 2.** ROI analysis results for production vs fixation cross and comprehension vs fixation cross contrasts, as well as the mean difference between them. Mean differences represent the group average of each subject’s mean vertex-wise activation difference. Asterisks indicate significant differences within a condition compared to the baseline (fixation cross) using a one-sample t-test: * *P <* 0.05, ** *P <* 0.01, *** *P <* 0.001. The p-value column indicates the significance of the production vs. comprehension comparison using a paired t-test. DMN (Overall) and FPN (Overall) show results when extracting data from the spatial union of their respective subnetwork masks.

| Network | Mean Prod<br>(vs Fix) | Mean Comp<br>(vs Fix) | Mean Diff.<br>(Comp-Prod) | $T_{29}$ | $P$ |
| --- | --- | --- | --- | --- | --- |
| DMN (Overall) | -0.076 | 0.857*** | 0.933 | 7.29 | < .001 |
| FPN (Overall) | 0.001 | 0.328* | 0.327 | 2.95 | .006 |
| LANG | 2.100*** | 3.286*** | 1.187 | 7.29 | < .001 |
| DN-A | -0.868*** | 0.643** | 1.511 | 9.54 | < .001 |
| DN-B | 0.710** | 1.211*** | 0.501 | 3.55 | .001 |
| FPN<sub>ext</sub> | 0.198 | 0.377* | 0.179 | 1.46 | .156 |
| FPN<sub>int</sub> | -0.227 | 0.333* | 0.560 | 3.93 | < .001 |

To compare conversational systems, we contrasted them directly to each other. The contrast **production > comprehension** revealed activation in 17 clusters (Fig. 3, Supplementary Table 3): (1, 3, 10, 11) bilateral pre- and postcentral gyri; (2, 8, 14-16) cerebellum; (4) bilateral supplementary motor area; (5-7, 12-13) basal ganglia; (9, 17) left inferior frontal gyrus.

**Table 3.** Paired t-test results comparing DMN and FPN subnetwork involvement in production and comprehension. Mean differences represent the direct subtraction of the respective subnetwork beta estimates (which were extracted from the production vs fixation and comprehension vs fixation contrasts).

| Condition | Comparison | Mean Diff. | $T_{29}$ | $P$ |
| --- | --- | --- | --- | --- |
| Production | DN-B – DN-A | 1.311 | 12.40 | < .001 |
|  | FPN<sub>ext</sub> – FPN<sub>int</sub> | 0.215 | 2.63 | .014 |
| Comprehension | DN-B – DN-A | 0.511 | 5.60 | < .001 |
|  | FPN<sub>ext</sub> – FPN<sub>int</sub> | -0.045 | -0.75 | .460 |

**Figure 3.**
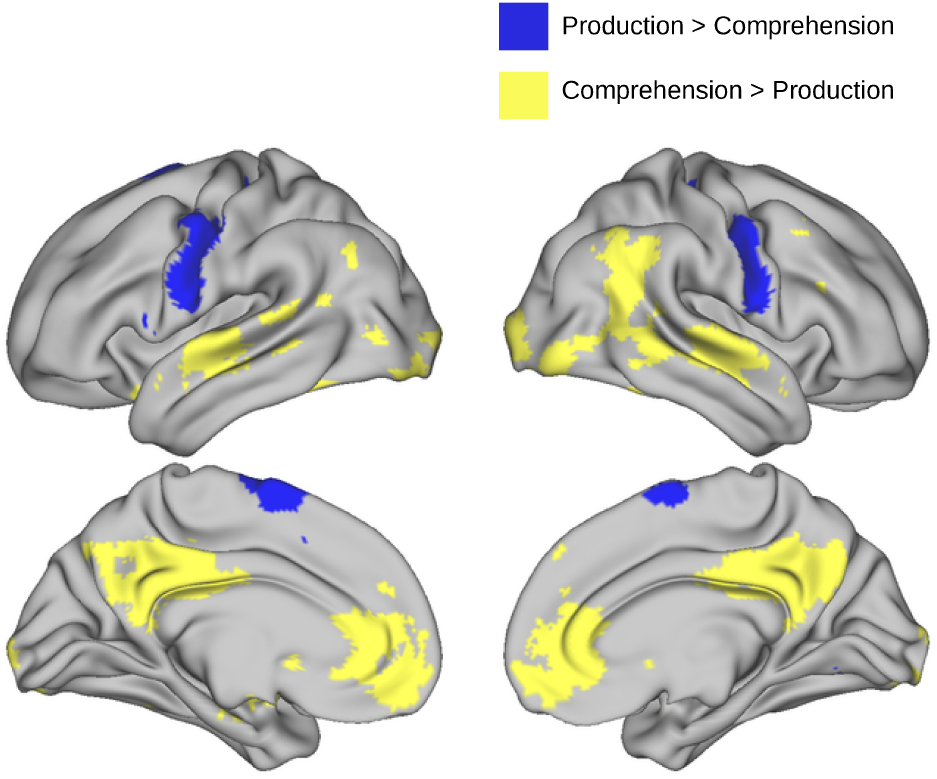
Whole-brain results for contrasting production vs comprehension and vice versa. Blue: areas more active for production over comprehension. Yellow: areas more active for comprehension over production. Cluster-forming threshold for these contrasts was set to *P* _FWE_ *<* 0.05.

The analysis of the frontal (LIFG) and temporal (LSTS) nodes activation within subject-specific language networks showed a stronger activation of the frontal node in production (mean diff. = 0.91, *T*_*29*_ = 3.99, *P<* 0.001), and stronger activation of the temporal node in comprehension (mean diff. = 1.54, *T*_*29*_ = 9.2, *P<* 0.001), see Table 4.

**Table 4.** Within the language network ROI analysis for the frontal and temporal nodes. ROIs are defined within subject-specific language network masks, intersected with LIFG and LSTS ROIs from Fedorenko et al. (2010) with 10 mm padding. Asterisks indicate significant differences within a condition compared to the baseline (fixation cross) using a one-sample t-test: *** *P <* 0.001. The p-value column indicates the significance of the production vs. comprehension comparison using a paired t-test.

| Language Network Node | Mean Prod<br>(vs Fix) | Mean Comp<br>(vs Fix) | Mean Diff.<br>(Comp-Prod) | $T_{29}$ | $P$ |
| --- | --- | --- | --- | --- | --- |
| Frontal (LIFG) | 2.412*** | 1.497*** | -0.915 | -3.99 | < .001 |
| Temporal (LSTS) | 2.592*** | 4.14*** | 1.549 | 9.2 | < .001 |

In the current study, we replicated previously found patterns of both similarities and asymmetries within the language network for conversational production and comprehension. This includes an asymmetry where the posterior IFG is preferentially involved for conversational production, while the superior and middle temporal gyrus are preferentially involved during conversational comprehension (see Fig. 2 and Table 4). Beyond the classical perisylvian language network, we replicate activity in a bilateral midline superior frontal area, as unique to production (see Fig. 3). Importantly, these asymmetries have thus now been shown to be stable: (1) across conversational lengths and (2) conversational topics, as the analyses of the Torubarova et al. (2025) and Rauchbauer et al. (2020) datasets demonstrate. We will discuss the connectivity (and neurodevelopmental) profile of the dorsal midline finding, in relation to the relevant networks, below. Using the paradigm with long conversations involving exchange of opinions, thus triggering live mentalizing, we have also shown involvement of the DMN during comprehension. This result is clear from Fig. 2, but was further specified using subject-specific functional network parcellation (MSHBM; Kong et al., 2019) of these networks (see Table 2). In this subject-specific network analysis, FPN networks were also observed to be active during comprehension.

These results align with the suggestion that production is the processing bottleneck during conversation (Levinson, 2024) due to increased pressure on the speaker to formulate a message considering syntactic structure as well as gestures. For comparison, the interleaved comprehension periods allow the listener to allocate more resources toward internally directed processes. Internally directed processes may, in other words, be observed during comprehension, while they are inherently about the conversation and thus tightly knit to upcoming turns as the speaker. Some examples of such processes are: (1) constructing the interlocutor’s opinion (Gumperz, 1996); (2) relating the interlocutor’s opinion to one’s own; (3) deciding whether to further the topic by persuading or alternatively exploring the interlocutor’s opinion (Guardiola and Bertrand, 2013); (4) exploring where to take the conversation next if the exchange is turning stale (Hoey, 2015); and (5) managing the conversation in other ways, e.g. managing turns (Levinson and Torreira, 2015), face/politeness (Brown et al., 1987) and common ground (Vanlangendonck et al., 2018). Note that e.g. (3)-(4) might be seen as early stages of the production process, although observed during comprehension.

As for FPN involvement during comprehension, this aligns with previous findings of its involvement in effortful context-dependent comprehension, which requires situational update (Smirnov et al., 2014; Yang et al., 2023), in contrast to passively processing words and sentences, where FPN engagement is absent (Diachek et al., 2020). It has also been suggested that FPN involvement in conversational comprehension can indicate the pressing demand for turn-taking management and planning one’s turn, while keeping track of the context, as in an ECoG study which showed frontolateral engagement during turn planning in an interactive language setting (Castellucci et al., 2022). In relation to our ethical dilemma scenario, an fNIRS study showed higher involvement of FPN for both production and comprehension during disagreement compared to agreement (Hirsch et al., 2021), which suggests that stance dis-/alignment further increases executive demands.

Taken together, stronger involvement of FPN and DMN during comprehension may indicate allocation of cognitive resources for a broader set of tasks such as active integration, management of alignment and turn-taking, to the comprehension time window, rather than the bottleneck production (Levinson, 2024) time window.

### Extralinguistic Networks: Theory of Mind and External Control as Essential

We will now further discuss this general pattern of activation in DMN and FPN during conversation by also considering their division into subnetworks. As already stated, conversational production has been suggested as a bottleneck for conversation, placing high priority on the rapid, uninterrupted execution of the planned turn as well as coordination with external turn- and feedback signals from the listener. This matches with our observation that FPN_ext_ activity was significantly higher than the suppressed FPN_int_ during production. At the same time, our results indicate that conversational production engages Theory of Mind (ToM) processes (Saxe and Kanwisher, 2003), as we observe DN-B activity (DiNicola and Buckner, 2021) for production. One possibility is that this activation may correspond to sociopragmatic processes related to taking the listener’ s perspective. Taken together, our results indicate that the taxing processes of formulation (language production areas), external control (FPN_ext_) and ToM (DN-B) may take priority over internally-oriented processes and thus suppress them. In our results, this corresponds to our observed suppression of DN-A and FPN_int_ during production.

At the same time, some internally oriented processing by interlocutors may be important to drive the conversation forward. Again, our results suggest that these processes are instead allocated to the comprehension time window. Conversational comprehension activates DN-A and FPN_int_ in our results, possibly as a consequence of the necessity, in this time window, to construct the situation model (DN-A) and provide domain general internal control of concurrent processes (FPN_int_). We note that both external and internal control processes cooperate during comprehension. External control (FPN_ext_) may be required e.g. to maintain focus on the speaker, importantly also including the visual input. This matches with our observation of the early visual cortex in comprehension > production. In accordance with this, external control (FPN_ext_) may be necessary e.g. to track rapid turn-taking cues, across modalities. Internal control (FPN_int_) may on the other hand exert top-down guidance to integrate the incoming narrative with internal schemas (via DN-A) and simultaneously actively plan one’ s upcoming turn (Levinson and Torreira, 2015).

Conversational comprehension also shows even stronger engagement of ToM processes, as indicated by stronger activation of DN-B during comprehension vs production. This is possibly a consequence of taking the speaker’s perspective to help derive the intended meaning, which might differ from the literal meaning. In summary, the findings when comparing subnetworks of DMN show that ToM (as indicated by activity in DN-B, which was stronger than activity in DN-A, across both comprehension and production) is an integral, continuous component of conversation, regardless of whether a person is currently speaking or listening, as interlocutors must constantly track their partner’ s epistemic state and intentions (Pickering and Garrod, 2013; Fedorenko et al., 2024).

### Further DMN Interpretations and Possible Divisions

In our further discussion on the DMN findings in relation to additional literature, we will again consider our results on DMN as a whole being involved in comprehension in relation to that literature, where the distinction of DN-A and DN-B is not always made. From a neurobiological perspective, there are several potential interpretations for DMN involvement during conversation, considering different suggested DMN functions. Building on the finding that the DMN encodes situation models and draws on such situation models to establish meaning (Fernandino and Binder, 2024; Thye et al., 2024b), such encoding and meaning processing can be predicted to happen specifically during comprehension, as this is when new information needs to be integrated. This aligns with our observations. Interestingly, our results also converge with those of Yamashita et al. (2025), who find several principal components linked to DMN for conversational comprehension, but none for production. For instance, Yamashita et al. (2025) found that four of the six principal components that explained most variance in the data were coupled with core DMN-activity. Their further investigation of the linguistic content of those components underscores the pragmatic function of the DMN during conversation, with a particular focus on comprehension, just as in our data. The four components were linked to material exemplified by, on the one hand self-referential utterances as (1) “I think” and “I would like to”, to utterances managing the conversation as in (2)-(3), where (2) had e.g. “uh”, “well”, “then”, and “you know what”, all often starting constituents and (3) with utterances requesting speech; and finally to other-mentalizing in DMN exemplified by (4) backchannels and laughter (Yamashita et al., 2025).

While these findings strongly link DMN activity to conversational pragmatics and situational modeling, it is important to emphasize that DMN involvement is not unique to the conversational setting. The DMN is also recruited during non-interactive narrative comprehension and production (Silbert et al., 2014; Stephens et al., 2010; AbdulSabur et al., 2014; Simony et al., 2016). Furthermore, cortical mappings of the semantic system demonstrate that regions overlapping with the DMN encode broad word-level meanings, particularly social ones, in a passive listening paradigm (Huth et al., 2016). Our observed DMN activity is not likely to be explained by task disengagement. When we manually annotate participant engagement utterance-by-utterance, participants are generally engaged in our paradigm (mean 4 on a 5-point scale, Torubarova et al., 2026) and DMN activity is not modulated by this annotated participant engagement (Ljung, 2026). Altogether, the DMN results for comprehension and production (in DN-B) show a broad and potentially varied involvement of the DMN during live mentalizing.

In this study, we conceptualize DMN as two parallel distributed systems (Braga and Buckner, 2017). However, the individual anatomical nodes comprising them are situated within distinct local topographies, with a robust body of literature discussing a functional gradient along the midline anterior-posterior DMN axis (e.g. Andrews-Hanna et al., 2014). We will now therefore also discuss those local findings in relation to conversation. It has been suggested (Stawarczyk et al., 2021) that the posterior midline DMN node (PCC) preferentially processes “other-related information and is involved in contextual information integration”. In contrast, the frontal node (mPFC) is specialized in more abstract self-referential processing, devoid of time and place, but with the important function of taking one’s desires into account. In the conversational context, this could correspond to increased conversational efforts focusing on (1) one’s interest in the interlocutor and management of the current conversation with the interlocutor, potentially centered on PCC vs (2) one’s thoughts on the topic, potentially centered on mPFC. This aligns with previous findings (Schieche et al., 2025; Bendtz et al., 2022), which found increased recruitment of the PCC, specifically during intention processing. In a dialogue comprehension paradigm, the study used indirect speech acts (containing a hidden face-saving speaker intention) vs direct speech acts (where the literal and intended meaning match) contrast. The sensitivity to this manipulation increased in the PCC along with increasing age through adolescence into adulthood (Schieche et al., 2025; Bendtz et al., 2022). In another recent study, narrative integration of sentences with high pragmatic and social content engaged the posterior mid-line DMN node (albeit with an emphasis on its more dorsal portion in precuneus, also together with the angular gyrus, Thye et al., 2024b) and yet another study on conversation activated this node specifically (Spatola and Chaminade, 2022; see Introduction).

### Network Considerations Around Posterior Dorsal Medial Frontal Finding

The current paper replicated previous findings of increased involvement of the dorsomedial prefrontal cortex (dmPFC), for the production vs fixation and production vs comprehension contrasts (Arvidsson et al., 2024b). The dmPFC was also modulated by lexical surprisal during conversational production (Arvidsson et al., 2024a). This is thus a stable finding related to conversational production, across shorter and longer conversations. While regions in the dmPFC, such as the preSMA, are generally linked to motor control, there is a growing body of fMRI evidence suggesting that these regions subserve language-specific processes (e.g., Chee et al., 1999; Moore-Parks et al., 2010; Carreiras et al., 2006) for instance when reading or listening to sentences (Wolna et al., 2026). Lesion and stimulation studies likewise show that disrupting the SMA can impair speech production (Jonas, 1981; Tremblay and Gracco, 2009), although recovery following unilateral lesions suggests functional compensation by the contralateral hemisphere (Riecker et al., 2010; Acioly et al., 2015). Consistent with these findings, network analyses have proposed that the preSMA and adjacent regions in the medial superior frontal gyrus forms part of an extended language network rather than the FPN/MD network (DiNicola and Buckner, 2021; Wolna et al., 2026; Ferstl et al., 2007), further supported by its functional connectivity profile (Du et al., 2024; Braga et al., 2020; Shain and Fedorenko, 2026). Our current and previous findings from conversation build on this evidence and further point to the possibility that, in the conversational context, the language processes localized in the dmPFC are more heavily recruited during production than comprehension.

While there are many possible explanations for such an asymmetry, we have previously suggested one such possible account: that socio-pragmatic (more posterior) dorsal medial prefrontal activations are driven by specialization to the conversational context. This observation aligns with a pattern wherein anterior regions handle abstract processes, while posterior areas are more engaged in context-specific functions along the frontal lobe’s anterior–posterior axis (Uddén and Bahlmann, 2012). Furthermore, previous developmental work demonstrates that as adolescents acquire greater pragmatic and conversational expertise, they increasingly recruit the posterior dorsal prefrontal cortex, in addition to more generalized ToM areas within the anterior and ventral sections of the medial prefrontal cortex (Forbes Schieche and Uddén, 2025). While this suggestion was made for both production and comprehension processes and the developmental data are based on pragmatic inference made during comprehension, the current data suggest revisiting the hypothesis, focusing on conversational production.

### Relation of Current Findings to Previous Large-Scale Models

Other large-scale models of cognitive neuroscience of language, such as Hickok (2022), also treat production vs comprehension differences in the language network. In contrast to Fedorenko et al. (2024), Hickock presents the language network as mainly a production network, but where some nodes are also relevant for comprehension. Our comprehension results converge with the posterior temporal and temporo-parietal regions that Hickok (2022) proposes as participating not only in production, but also in comprehension, including bilateral superior and middle temporal gyri and the angular gyrus. Our results consistently show superior temporal areas to be preferentially (or sometimes uniquely) activated by comprehension (Arvidsson et al., 2024b; replicated in Goldstein et al., 2025). This raises the question of what role this region plays during production in conversational context. One possibility is that this activity reflects an efference copy, i.e., an auditory prediction of one’s own speech which supports self-monitoring during speaking (as suggested by Zada et al., 2024).

On the other hand, Fedorenko et al. (2024) laudably encompass dynamics between extra-linguistic networks and the language network. While our results (together with Goldstein et al., 2025) support the general directions in this model, they also suggest a more complex specialization within the language and extra-linguistic networks, for comprehension vs production. An important example is the IFG, which shows earlier activity relative to the STG in production, but later activity in comprehension (Goldstein et al., 2025; Uddén et al., 2022, see also Giglio et al., 2024). As a whole, in the subject-specific network analysis, the language network was more active during comprehension than production. This finding has not yet been incorporated in the large-scale models in cognitive neuroscience of language. Whole-network results that come out of a whole-brain analysis or subject-specific network analysis (such as those in Table 2) are important for interpreting large-scale networks as characterizing coherent patterns of task-relevant functional connectivity, rather than isolated regional responses. Note that the FPN results are considered less robust compared to the DMN results, as they mainly rely on the subject-specific network analysis.

## Limitations

We interpret that DMN response in the current results as potentially being a consequence of the more intense live mentalizing in the current paradigm (due to longer interactions involving discussion of opinions on dilemmas). However, increased power may also explain the observation of DMN in the current paper, but not in Arvidsson et al. (2024b). Arguing against this is the complete absence (even subthreshold) of DMN activity in that study, as the current DMN activations are strong.

While face-to-face interaction and conversation entail a spectrum of interaction forms, where fewer constraints are more ecologically valid (Kendrick and Holler, 2024), our current paradigm is still suitable for studying conversation with fMRI. The method has high spatial resolution and good coverage of medial structures (compared to, e.g., ECoG), ideal for our research questions. While it is true that fast processes cannot be directly targeted using fMRI, many fast processes in the same area, on this time scale, will add up to a metabolic consequence, visible with fMRI (Yeşilyurt et al., 2008).

We have framed this study as using a naturalistic fMRI paradigm, and while we believe the naturalistic turn will stay, note that such paradigms come with their own issues (Leipold et al., 2024). Neural results are likely to depend on the topic, interlocutor relation, and content/differentiation of speech acts, all parameters for further study. Moreover, while it was crucial for our intended manipulation to use a confederate instead of a naive interlocutor, it is important to acknowledge that confederates may hinder the naturalness of an interaction even when the confederate’s task is to behave naturally (Kuhlen and Brennan, 2013).

Our approach robustly isolates the foundational differences in the neural infrastructure of production and comprehension, but it is limited as we don’t consider signal changes within blocks of production and comprehension. While this method allows us to handle the high variability within live, unscripted conversations, further research should utilize the dynamic features that a natural conversation provides. Alternative approaches may include inter-regional connectivity to assess the synchronization between language and extra-linguistic networks (Paunov et al., 2019), leveraging the variability in the conversational content (Thye et al., 2024a, 2025), and/or integrating multimodal features in order to optimize a neural *foundation model*, i.e. a model to predict ongoing neural responses across the brain from a rich set of annotated or automatically derived latent features describing the naturalistic stimuli or interactive situation (Santana et al., 2026, under review; d’Ascoli et al., 2026).

## Conclusion

By revisiting classical questions of language processing, but now with conversational data, we have established robust asymmetries between production and comprehension, including LIFG preferences for production and S/MTG preferences for comprehension. We have further delineated an area in the posterior dorsal medial pre-frontal cortex, with unique activation for conversational production. Finally, we have also shown robust default mode network involvement and additional involvement of the frontoparietal network for control, during conversational comprehension, when considering one’s own and others’ opinions in longer exchanges (i.e. live mentalizing). A subject-specific network analysis showed that the subnetwork DN-B was more strongly activated than DN-A, across both comprehension and production. The results indicate that there are strong pressures for language formulation, external directed control, and Theory of Mind (suppressing internally oriented processes) during conversation production. Comprehension may thus allow for more internally oriented processing, situation-model construction, as well as even stronger Theory of Mind engagement that support understanding and help prepare responses, however with Theory of Mind processes remaining continuously active throughout conversation. Combined, our results substantially further the understanding of the involvement of the linguistic and extralinguistic networks during conversational production and comprehension.

## Ethical Approval

Swedish Ethical Review Authority (Dnr: 2021-06225-01) gave approval for the original study. Informed consent was obtained from all participants. No new data collected in the current paper.

## Data Availability

Dataset available at https://openneuro.org/datasets/ds004996.

1 https://www.nitrc.org/projects/art_repair/

2 https://github.com/ThomasYeoLab/CBIG/tree/master/stable_-projects/brain_parcellation/Kong2019_MSHBM

