## Supplementary Material for "Conversation in fMRI: DMN Activates During Live Mentalizing"

**S-1: Event Duration and Motion Statistics**

| **Event Type** | **Eng. Level** | **N of Events** | **Event Duration (s)** | **Mean FD (mm)** |
| --- | --- | --- | --- | --- |
| **Production** | High | 130.3 (SD = 28.7) | 1.98 (SD = 1.56) | 0.29 (SD = 0.27) |
|  | Medium | 155.8 (SD = 27.3) | 2.03 (SD = 1.51) | 0.28 (SD = 0.25) |
|  | Low | 144.0 (SD = 25.7) | 2.04 (SD = 1.51) | 0.28 (SD = 0.27) |
| **Comprehension** | High | 135.9 (SD = 31.2) | 2.17 (SD = 2.61) | 0.23 (SD = 0.22) |
|  | Medium | 113.8 (SD = 20.2) | 1.70 (SD = 2.39) | 0.23 (SD = 0.23) |
|  | Low | 61.9 (SD = 17.7) | 2.50 (SD = 2.39) | 0.18 (SD = 0.17) |
| **Silence** | High | 131.1 (SD = 29.9) | 0.76 (SD = 0.71) | 0.24 (SD = 0.22) |
|  | Medium | 155.0 (SD = 29.8) | 0.85 (SD = 0.69) | 0.25 (SD = 0.22) |
|  | Low | 156.2 (SD = 24.6) | 1.08 (SD = 1.06) | 0.25 (SD = 0.23) |

**Supplementary Table 1.** Descriptive statistics of the data per subject for each engagement condition: average number and duration of events, mean framewise displacement (FD) and percentage of scans with excessive motion (FD > 1mm).

To evaluate possible effect of motion on the whole-brain results, we ran an additional post-hoc analysis, splitting the subjects into low and high motion groups based on mean FD. We estimated two second-level models with two-sample t-tests (assuming independence and unequal variance) for production vs fixation and comprehension vs fixation first-level contrasts. There were no significant clusters between low and high motion groups for either of the contrasts (also at a lenient cluster-forming threshold *P_uncorr_* < 0.005). We then compared contrast estimates for the subject-specific language networks across the low and high motion groups. There was no significant difference in language network involvement dependent on motion (production vs fixation *T_27.59_* = -0.53, *P* = 0.59; comprehension vs fixation *T_27.7_* = -0.88, *P* = 0.38).

**S-2: MSHBM Parcellation Validation**

To validate individual network parcellation conducted via MSHBM algorithm (Kong et al., 2019), we calculated a spatial overlap between two canonical atlases (Yeo et al., 2011; Du et al., 2014) and individual networks, taking the highest overlap as the closest network match. The mapping was consistent for all subjects. Specifically, the closest match for Yeo et al. (2011) and Du et al. (2014) atlases, respectively, was as follows: TempPar – LANG; Default A – DN-A; Default B – DN-B; Control A – FPN-A; Control B – FPN-B.

As follows from its name, Yeo et al. (2011) TempPar network covers temporal and parietal cortices and lacks frontal regions, crucial for a candidate language network. The spatial overlap score between TempPar and LANG was 0.44. The average overlap of individual networks, however, showed a higher match to Du et al. (2014) LANG than Yeo et al. (2011) TempPar (M = 0.57). To assess the spatial distribution of this shift, we first created a probabilistic map of individual networks and subtracted Yeo et al. (2011) TempPar map, applying a threshold of 70% agreement (i.e. only keeping the vertices where >70% of the subjects expanded the TempPar network). This excess map included the left IFG and SFG, as well as bilateral STS and STG, with a strong left-lateralized preference. This procedure confirmed that the individually generated maps form reliable language network candidates. See consensus atlas of individually estimated networks in Figure S1.

To evaluate intra-subject network stability, we ran another MSHBM parcellation for every subject’s run separately. Mean intra-subject stability (run-to-run Dice overlap) across all the networks of interest = 0.92 (SD = 0.009). Mean inter-subject overlap across the relevant networks = 0.87 (SD = 0.01). Very stable results are likely due to a small amount of data, which causes the Bayesian model to rely more heavily on the group prior; however, for every network, intra-subject stability across runs was greater than inter-subject overlap (*P* < 0.001), in accordance with Kong et al. (2019).

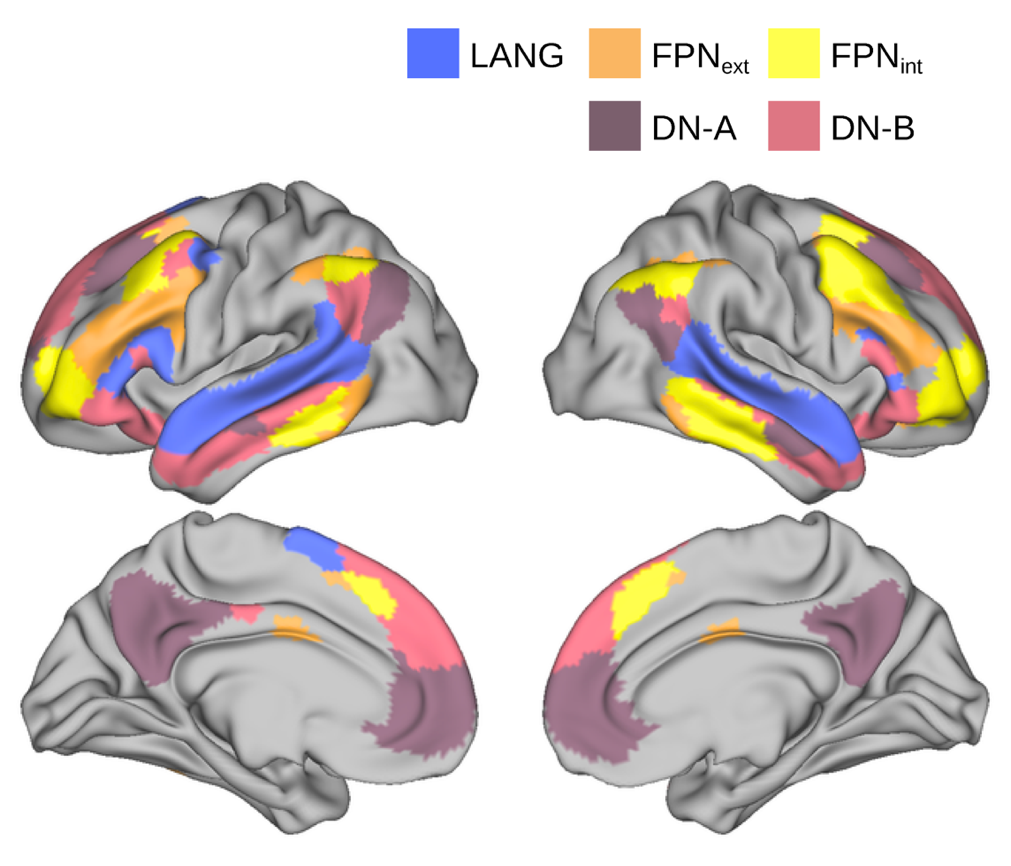

**Figure S1.** Consensus atlas of the individualized functional networks in our data. Each vertex is assigned to the network with the highest spatial probability across subjects, thresholded to display only vertices with >20% overlap across subjects.

**S-3: List of Abbreviations for Brain Areas**

ACC - anterior cingulate cortex

IFG - inferior frontal gyrus

IOG - inferior occipital gyrus

ITG - inferior temporal gyrus

MFG - middle frontal gyrus

MOG - middle occipital gyrus

MTG - middle temporal gyrus

MCC - midcingulate cortex

OFC - orbitofrontal cortex

SFG - superior frontal gyrus

SMA - supplementary motor area

SOG - superior occipital gyrus

STG - superior temporal gyrus

Hipp. – hippocampus

Parahipp. – parahippocampal gyrus

**S-4: fMRI Results with MNI Coordinates**

**Supplementary Table 2.** Significant activations in the production > fixation, comprehension > fixation contrasts and their conjunction. In this and the following table, for each cluster, MNI coordinates of the cluster’s local maxima are given together with cluster size.

| **Anatomical region** | **MNI local maxima** | | | **cluster** | | **voxel** | |
| --- | --- | --- | --- | --- | --- | --- | --- |
|  | **x** | **y** | **z** | **size** | ***P_FWE_*** | ***T*-value** | ***P_FWE_*** |
| **1) Production > Fixation Cross (cluster-forming threshold *P_FWE_* < 0.05)** | | | | | | | |
| R precentral, R postcentral | -52 | -12 | 40 | 932 | .000 | 16.24 | .000 |
| L precentral, L postcentral | 52 | -14 | 38 | 1504 | .000 | 15.47 | .000 |
| L/R cerebellum | 14 | -62 | -16 | 3258 | .000 | 14.72 | .000 |
| L/R SMA | -4 | 0 | 64 | 703 | .000 | 12.65 | .000 |
| L MTG, L STG | -64 | -26 | 6 | 847 | .000 | 11.38 | .000 |
| R STG | 62 | -4 | 2 | 276 | .000 | 9.95 | .000 |
| L IFG | -46 | 14 | 14 | 137 | .000 | 9.56 | .000 |
| L basal ganglia | -8 | -22 | -12 | 43 | .000 | 9.36 | .000 |
| L basal ganglia | -22 | -12 | -6 | 27 | .000 | 9.12 | .000 |
| L basal ganglia | -14 | -18 | 4 | 37 | .000 | 8.54 | .000 |
| R cerebellum | 26 | -78 | -40 | 113 | .000 | 8.17 | .001 |
| L superior temporal pole | -54 | 14 | -18 | 77 | .000 | 7.84 | .001 |
| L lingual, L MOG, L calcarine | -14 | -90 | -10 | 25 | .000 | 7.53 | .001 |
| L SMA | -4 | 14 | 48 | 21 | .000 | 7.42 | .004 |
| L cerebellum | -14 | -64 | -50 | 32 | .000 | 7.22 | .006 |
| R calcarine | 18 | -94 | 2 | 37 | .000 | 6.85 | .014 |
| **2) Comprehension > Fixation Cross (cluster-forming threshold *P_FWE_* < 0.05)** | | | | | | | |
| L MTG, L STG | -58 | -12 | 6 | 3950 | .000 | 15.7 | .000 |
| R STG, R MTG | 62 | -6 | 0 | 2308 | .000 | 15.11 | .000 |
| L/R calcarine | -34 | -84 | -18 | 1160 | .000 | 11.17 | .000 |
| L cerebellum | -26 | -82 | -36 | 249 | .000 | 10.79 | .000 |
| R cerebellum | 18 | -80 | -36 | 599 | .000 | 10.45 | .000 |
| L fusiform, L ITG | -42 | -46 | -18 | 157 | .000 | 9.12 | .000 |
| R IOG, R MTG | 50 | -78 | -4 | 330 | .000 | 8.95 | .000 |
| L SFG, R SFG | -6 | 54 | 42 | 321 | .000 | 8.73 | .000 |
| R cerebellum | 14 | -62 | -46 | 14 | .000 | 8.46 | .000 |
| L thalamus, L lingual | -10 | -32 | -2 | 45 | .000 | 8.43 | .000 |
| L thalamus | 12 | -28 | -2 | 30 | .000 | 8.38 | .000 |
| R fusiform | 42 | -50 | -22 | 24 | .000 | 7.54 | .003 |
| L IFG | -44 | 16 | 30 | 74 | .000 | 7.48 | .003 |
| R cerebellum | 8 | -52 | -42 | 33 | .000 | 7.41 | .004 |
| L IFG | -50 | 30 | -8 | 13 | .000 | 7.17 | .008 |
| L MFG, L precentral | -48 | 4 | 52 | 19 | .000 | 7.11 | .009 |
| L cerebellum | -4 | -56 | -42 | 15 | .000 | 7.05 | .015 |
| L MOG | -48 | -76 | 6 | 15 | .000 | 6.9 | .015 |
| **3) Conjunction Analysis (cluster-forming threshold *P_FWE_* < 0.05)** | | | | | | | |
| R STG, R Heschl’s | 56 | -16 | 6 | 464 | .000 | 8.99 | .000 |
| L superior temporal pole | -52 | 14 | -18 | 299 | .000 | 8.85 | .000 |
| L MTG, L STG | -60 | -12 | 6 | 1242 | .000 | 8.32 | .000 |
| R cerebellum | 26 | -82 | -38 | 176 | .000 | 8.2 | .000 |
| R cerebellum | 18 | -72 | -26 | 63 | .000 | 7.75 | .000 |
| L/R calcarine | 16 | -92 | 0 | 328 | .000 | 7.47 | .000 |
| L cerebellum, L fusiform | -40 | -72 | -20 | 24 | .000 | 6.57 | .002 |
| R fusiform, R cerebellum | 42 | -70 | -20 | 16 | .001 | 6.55 | .002 |
| L IFG | -52 | 32 | -6 | 30 | .000 | 6.5 | .003 |
| R middle/superior temporal pole | 48 | 16 | -24 | 29 | .000 | 6.49 | .003 |
| L SFG | -10 | 46 | 44 | 15 | .002 | 6.35 | .005 |
| L SFG | -44 | 8 | 60 | 50 | .001 | 6.23 | .007 |
| L IFG | -54 | 18 | 18 | 17 | .001 | 6.12 | .01 |
| R cerebellum | 4 | -56 | -56 | 15 | .002 | 6.11 | .01 |
| L IFG | -52 | 24 | 12 | 25 | .000 | 5.95 | .017 |

**Supplementary Table 3.** Significant activations in production > comprehension and comprehension > production contrasts.

| **Anatomical region** | **MNI local maxima** | | | **cluster** | | **voxel** | |
| --- | --- | --- | --- | --- | --- | --- | --- |
|  | **x** | **y** | **z** | **size** | ***P_FWE_*** | ***T*-value** | ***P_FWE_*** |
| **1) Production > Comprehension (cluster-forming threshold *P_FWE_* < 0.05)** | | | | | | | |
| L postcentral, L precentral | -50 | -12 | 40 | 1580 | .000 | 19.0 | .000 |
| L/R cerebellum | 14 | -58 | -22 | 2460 | .000 | 18.11 | .000 |
| R precentral, R postcentral | 46 | -10 | 34 | 1187 | .000 | 16.3 | .000 |
| L/R SMA | -2 | 4 | 72 | 852 | .000 | 13.1 | .000 |
| L basal ganglia | -16 | 2 | 26 | 58 | .000 | 9.46 | .000 |
| L basal ganglia | -24 | -12 | -2 | 102 | .000 | 9.43 | .000 |
| L basal ganglia | -30 | -68 | 10 | 28 | .000 | 8.42 | .000 |
| R cerebellum | 28 | -60 | -48 | 39 | .000 | 8.37 | .000 |
| L IFG | -40 | 14 | 14 | 24 | .000 | 8.32 | .001 |
| R precentral, R postcentral | 22 | -30 | 58 | 61 | .000 | 8.18 | .001 |
| L postcentral, L precentral | -18 | -30 | 60 | 67 | .000 | 8.12 | .001 |
| L basal ganglia | -34 | -54 | 0 | 18 | .000 | 7.72 | .002 |
| R basal ganglia | 24 | -10 | -4 | 21 | .000 | 7.68 | .002 |
| R cerebellum | 12 | -66 | -42 | 39 | .000 | 7.51 | .003 |
| L cerebellum | -14 | -66 | -40 | 18 | .000 | 7.29 | .005 |
| L cerebellum | -6 | -42 | -22 | 18 | .000 | 7.03 | .009 |
| L IFG | -50 | 14 | 8 | 28 | .000 | 6.89 | .012 |
| **2) Comprehension > Production (cluster-forming threshold *P_FWE_* < 0.05)** | | | | | | | |
| L/R precuneus, L/R MCC | -2 | -32 | 30 | 2638 | .000 | 14.11 | .000 |
| L MTG, L STG | -56 | -14 | -8 | 872 | .000 | 13.93 | .000 |
| R MTG, R STG, R angular | 60 | -12 | -6 | 3869 | .000 | 13.9 | .000 |
| L/R ACC, L SFG, L/R MFG | 0 | 46 | 10 | 2353 | .000 | 12.3 | .000 |
| L hipp, L amygdala, L parahipp. | -16 | -2 | -18 | 295 | .000 | 10.71 | .000 |
| L MTG, L STG | -66 | -42 | 12 | 334 | .000 | 10.08 | .000 |
| L basal ganglia | -4 | 10 | -2 | 50 | .000 | 9.95 | .000 |
| L fusiform, L ITG | -44 | -52 | -20 | 145 | .000 | 9.71 | .000 |
| R fusiform, R ITG | 42 | -44 | -14 | 74 | .000 | 9.23 | .000 |
| L cerebellum | -12 | -84 | -28 | 326 | .000 | 9.06 | .000 |
| L angular, L MTG, L MOG | -56 | -62 | 42 | 656 | .000 | 9.02 | .000 |
| L MOG, L IOG, L calcarine | -30 | -100 | 2 | 581 | .000 | 8.91 | .000 |
| R lingual, R IOG, R calcarine | 22 | -88 | -6 | 150 | .000 | 8.68 | .000 |
| L OFC, L insula, L superior temporal pole | -30 | 16 | -20 | 38 | .000 | 8.17 | .001 |
| R superior/middle temporal pole | 58 | 10 | -18 | 22 | .000 | 7.76 | .002 |
| R parahipp., R hipp. | 18 | -4 | -18 | 15 | .000 | 7.64 | .002 |
| R MFG | 42 | 20 | 52 | 35 | .000 | 7.39 | .002 |
| R OFC, R superior temporal pole | 30 | 8 | -20 | 17 | .000 | 7.32 | .004 |
| L/R SFG | 2 | 46 | 42 | 21 | .000 | 7.29 | .005 |
| R IFG | 54 | 32 | 24 | 26 | .000 | 7.12 | .007 |
